# A neural correlate of learning fails to predict foraging efficiency in *Bombus terrestris*

**DOI:** 10.1101/2024.01.23.576659

**Authors:** Grégoire Pasquier, Christopher D. Pull, Swidbert R. Ott, Ellouise Leadbeater

## Abstract

The mushroom bodies (MB) are integrative structures in the insect brain that, in social bees, contribute to both visual and olfactory learning. Changes in the density of presynaptic boutons (or microglomeruli) within the calyx region of the MB have been repeatedly linked to various aspects of foraging, including forms of learning that are thought to be key to support foraging efficiency. Here we set out to directly test the relationship between foraging efficiency and microglomerulus density in a bumblebee model, *Bombus terrestris.* We found no evidence that microglomerulus density predicted real-world foraging performance, nor any relationship with foraging experience. Instead, our data suggest a potential non-linear relationship between individual age, independent of foraging experience, and microglomerulus density in the lip region of the calyx, which is associated with olfactory processing. Our findings suggest that in real-world scenarios there is no simple direct relationship between microglomerular density, learning ability and foraging efficiency in bumblebees, highlighting the gap in knowledge regarding the relationships between learning abilities, neuroanatomy and foraging efficiency.

## Introduction

The workers of social bees face the challenge of collecting many small nectar and pollen rewards across a relatively vast foraging range. The ability to learn and remember floral characteristics that predict reward, alongside the locations at which rewarding patches have been found and the current reward levels available within them, is thought to be integral to the efficient fulfilment of this task (Chittka and Thomson 2001; Klein et al. 2017). Accordingly, a body of work has shown that the microstructure of the mushroom bodies (MB), which are integrative neural structures that are associated with learning and memory abilities, is plastic and reflects aspects of engagement in foraging tasks (Withers et al. 1993; Durst et al. 1994; Ismail et al. 2006; Groh et al. 2012; Scholl et al. 2014; Muenz et al. 2015; Cabirol et al. 2018). Intrinsic neurons in the MB, the Kenyon cells, connect to dendrites coming from sensory neurons to form synaptic boutons also known as microglomeruli (MG) (Groh and Rössler 2011). Given that neurogenesis does not take place in the MBs of adult insects (Fahrbach et al. 1995), it is these structures that have been the focus of studies linking MB structural variation to learning performance in bees (Hourcade et al. 2010; Van Nest et al. 2017; Li et al. 2017). Each MG is a synaptic complex that contains a central cholinergic synaptic bouton projecting from the antennal or optic lobes, surrounded by smaller GABAergic or octopaminergic boutons (Frambach et al. 2004).

In honeybees (*Apis mellifera*), where workers exhibit temporal polytheism, the onset of foraging coincides with a decrease in the density of MG in both the collar and the lip region of the mushroom bodies, which house projections from the antennal and olfactory lobes, respectively (Groh et al. 2012). This pruning of projection neuron boutons is not age-dependent but triggered by foraging itself, through exposure to light (Scholl et al. 2014). It is accompanied by an increase of Kenyon cell dendrites as well as overall volume of the mushroom bodies (Withers et al. 1993; Farris et al. 2001) and has been hypothesised to prime the brain for learning about floral rewards (Withers et al. 1993; Farris et al. 2001; Cabirol et al. 2018). Computational modelling indeed suggests that sparser coding of information within the MBs is optimal for updating learnt information, allowing for the formation of new associations to guide decisions and thus maximise foraging efficiency (Cabirol et al. 2018). MG density subsequently increases again as bees learn about their environment and accumulate foraging experience (Cabirol et al. 2018). Accordingly, learning events such as formation of an associative memory result in an increase in MG density in honeybees (Hourcade et al. 2010), and potentially also in bumblebees (*Bombus* spp,; Li et al. 2017).

These findings, and related studies in ants (Stieb et al. 2010), suggest that mushroom body plasticity in social insects may respond to both the need for, and the process of, learning and memory retrieval, in order to support efficient foraging (Fahrbach and Van Nest 2016; Cabirol et al. 2018). Indeed, both longer- and shorter-term memory performance (as captured through laboratory assays) have been found to correlate with foraging efficiency in bumblebees that forage in the real world (although these effects can be specific to particular environments; Raine and Chittka 2008; Pull et al. 2022). However, no study has yet explored whether this relationship might be driven by neural structure.

Here, we directly test the relationship between foraging efficiency, foraging experience and MG density in the bumblebee *Bombus terrestris audax*. Unlike honeybees, bumblebees do not exhibit age-based polyethism and, accordingly, MG density reduction occurs earlier in the life-cycle than it does in honeybees, in preparation for the commencement of foraging within 2-3 days of emergence from the pupal stage (Kraft et al. 2019). We hypothesize that, when sampled mid-lifespan, MG density may be greater in those bees that have engaged in more foraging trips, and in those that forage more efficiently.

## Methods

### Overview

Following a staggered design (Figure 1), bees of known age with no previous foraging experience were fitted with radio-frequency identification chips (RFID) and allowed to forage freely for 8 days (= ∼ 40% of foraging lifespan) using a hole-in-the-wall set-up (Raine and Chittka 2008; Evans et al. 2017; Pull et al. 2022). All bees originated from laboratory-reared, commercially acquired colonies raised under identical conditions. Foraging efficiency was recorded for each individual as mass of nectar collected per minute, over multiple recorded foraging trips, before quantification of MG density through synapsin-based immunostaining of both the lip and collar regions.

**Figure 1:**
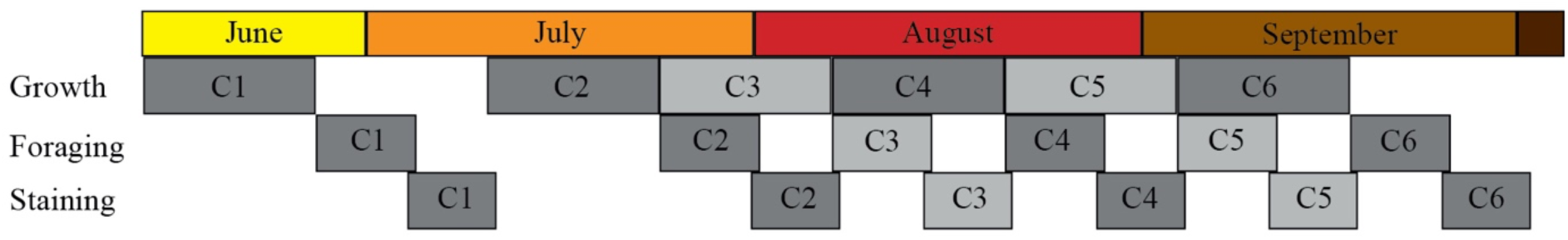
Timeline of the experiment. Each colony (C1-6, n = 6 colonies) went through a growth phase, during which the workers that were later sampled emerged and were tagged. At the end of the foraging period, foraging bees were sampled and their brain tissues were processed following a standard immunostaining protocol. The time gap between C1 and C2 is due to a colony producing only 3 workers by the end of the growth phase, which prompted its removal from the study.

### Growth phase

Within each colony, we created a cohort of tagged and RFID-chipped bees that were of known age and had never foraged before. Six *Bombus terrestris audax* colonies in the very early phase of worker number expansion (∼1 week following initial worker emergence) were obtained from a commercial supplier (Agralan, UK). Upon receipt of a colony, we standardized the colony size to 20 workers, which we tagged by supergluing (cyanoacrylate, Loctite) plastic number-discs on the dorsal thorax to identify them as having emerged pre-arrival. The queen and the brood were left undisturbed. Individual age could only be defined for bees that emerged after receipt of the colony from the supplier. To maximise the number of individuals of known age, the colony was allowed to grow in size in the laboratory for two weeks. During this growth phase (Figure 1), the focal colony was kept in a dark room at 24°C with sugar solution (35% w/w) provided *ad libitum* and pollen added three times per week. Three times a week, newly emerged bees were individually tagged (as above) under red light (that bees cannot see) with an RFID chip and a plastic numbered disc on the dorsal part of their thorax (the RFID chip was placed underneath the numbered disc).

### Ethical Note

The experiment described here followed the ASAB/ABS Guidelines for the use of animals in Research and did not require any license or permits in the United Kingdom. The colony boxes provided a dark environment which aimed to replicate the underground conditions *Bombus terrestris* colonies prefer in nature. The tags glued on the thorax of the bees did not prevent normal flight behaviour. The colonies were euthanised by freezing them after the experiment was completed.

### Foraging phase

During this phase, tagged and chipped bees were provided with unlimited outdoor access to accumulate foraging experience. To that end, we removed access to the sugar solution from colonies two days before the end of the growth phase, ensuring that colonies were motivated to forage while still having sufficient stores to avoid starvation. On the first day of the foraging phase, colonies were rehoused under red light into a clean nest box made of grey Perspex (28×16×10.5 cm^3^) that opened to a clear Perspex tunnel fitted onto a precision scale (Ohaus Advanced Portable Balance Scout STX) and an RFID reader system (MAJA Bundle Bee Identification System iID2000, ISO15693 optimized, Micro-Sensys GmbH). The tunnel had a false bottom so that bees ran directly across the pan of scales to record their weight using the weight-averaging function. The tunnel was in turn connected to a clear plastic tube giving access to outdoors through a hole cut in the window of the laboratory. During the foraging phase, the colony was allowed to forage freely on and around our university campus (Egham, Surrey, UK), which comprises mixed woodland, parkland and planted ornamental gardens. The campus and private gardens in the surrounding area provided flowering plants to the colonies throughout the experiment and no additional sugar solution was required to feed the colonies during this phase. Pollen was provided *ad libitum* throughout to reduce pollen foraging.

Colonies were monitored by a single observer for 6 hours (between 0815 and 1630 GMT) on five different days of the foraging phase, during which time the mass on exit and entry of all foraging workers was recorded (mean of three measurements using the scale’s 2-second weight averaging function taken during each tunnel crossing, i.e., each every entry/exit event, which minimises noise generated by the bees movement), alongside the time taken for each foraging trip. Foraging efficiency was calculated for each foraging trip as the weight difference between exit and entry, divided by trip duration.

### Brain fixation and dissection

At the end of the Foraging phase, all foragers were sampled upon returning from a foraging trip. The subjects were chilled on ice for 10 minutes and decapitated using dissecting scissors. The heads were pinned on a dissection plate and submerged in 0.1 M HEPES-buffered saline (HBS). We cut a large square window in the front of the head capsule and removed all of the air sacs around the brain that were accessible through the window, thus exposing the frontal surface of the brain. The heads were then transferred into ice-cold 4% formaldehyde in 0.1 M phosphate-buffered saline (PBS; pH = 7.4) and fixed overnight at 4°C on an orbital shaker, washed in HBS twice for 5 minutes, pinned again on a dissection plate and finally submerged in HBS. The compound eyes, ocelli and the remaining air sac membranes at the back of the brain were removed. Finally, the last remaining anchor points of the brain in the head capsule were severed by an incision under the antennal lobes. The free-floating brains were then washed in 0.1 M PBS twice for 5 minutes at room temperature on an orbital shaker.

### Cutting frozen sections

The fixed brains were cryoprotected using a graded series of sucrose (10%, 20% and 30%) in 0.1 M PB with 0.005% sodium azide (NaN_3_). Each step of the graded series lasted 1 h at room temperature, ensuring that the brains had sunk to the bottom of the vial. The brains were stored in 30% sucrose/phosphate buffer (PB) at 4°C for two days, then embedded in aqueous 20% gelatine solution in stainless steel moulds, oriented in the embedding medium with the frontal side facing down. The moulds were placed on dry ice until the gelatine was uniformly frozen. The gelatine block was fitted to a chuck and left in the cryostat chamber for 15 min to give the object time to equilibrate its temperature with the chamber. Sections were cut at -18°C to a thickness of 30 µm, arranged on Superfrost Plus™ Adhesion Slides (Thermo-Fisher) and stored overnight at 4°C to let them thaw and dry.

### Antibody incubation and mounting

The slides were placed on a hot plate for approximately 5 s to melt the gelatine before being rehydrated for 10 minutes in 0.1 M PB in a Coplin jar. All subsequent washes were done in 0.1 M PBS with 0.2% Triton X-100 (PBSTx). Slides were rinsed in PBSTx and pre-incubated with 5% normal goat serum in PBSTx (NGS/ PBSTx) for 45 minutes. Slide edges were framed with a hydrophobic pen (Advanced PAP Pen, Sigma-Aldrich) around the edges, covered with 200 µl of anti-synapsin primary antibody 1:50 in NGS/PBSTx (3C11 monoclonal mouse anti-SYNORF1, Klagges et al. 1996; Developmental Studies Hybridoma Bank, University of Iowa, USA), for 2.5 hours in a moisture chamber.

The slides were briefly rinsed in PBSTx, washed 4 times in PBSTx for 10 minutes, and then incubated with Cy3-conjugated affinity-purified goat anti-mouse IgG (H+L) polyclonal antibody (1:100; ThermoFisher, cat.no. A10521) and Alexa 488-conjugated phalloidin (1:200; ThermoFisher, cat.no. A12379) in NGS/ PBSTx. Incubation was as for the primary antibody, but protected from direct light with an aluminium foil cover. After 1.5 h, slides were rinsed in PBSTx before being washed 4 times in PBSTx for 10 minutes and mounted in 90% glycerol/PB containing 3% n-propyl gallate as anti-fading agent.

### Confocal microscopy and microglomerulus density measurement

Stacks of confocal images of the mushroom bodies were captured on an Olympus FV-10 laser scanning confocal microscope (Olympus Corporation) with a 60x oil immersion objective (Olympus UPLSAPO 60xo, NA = 1.35) and z-spacing of 0.41 µm. For each subject, we chose 4 contiguous physical 30 µm sections. We randomly assigned one of the four calices to each physical section so that each one of the left lateral, the left medial, the right medial and the right lateral calyx were scanned once per individual. The stacks were analysed with the software ImageJ (Rasband 2010). We used a random offset grid to place three cubes (8.2 `×` 8.2 `×` 8.2 μm^3^) in the dense collar region and three cubes in the lip region of the calyx (Fig. 2). We then manually counted the presynaptic boutons contained in the cubes to quantify their density.

**Figure 2:**
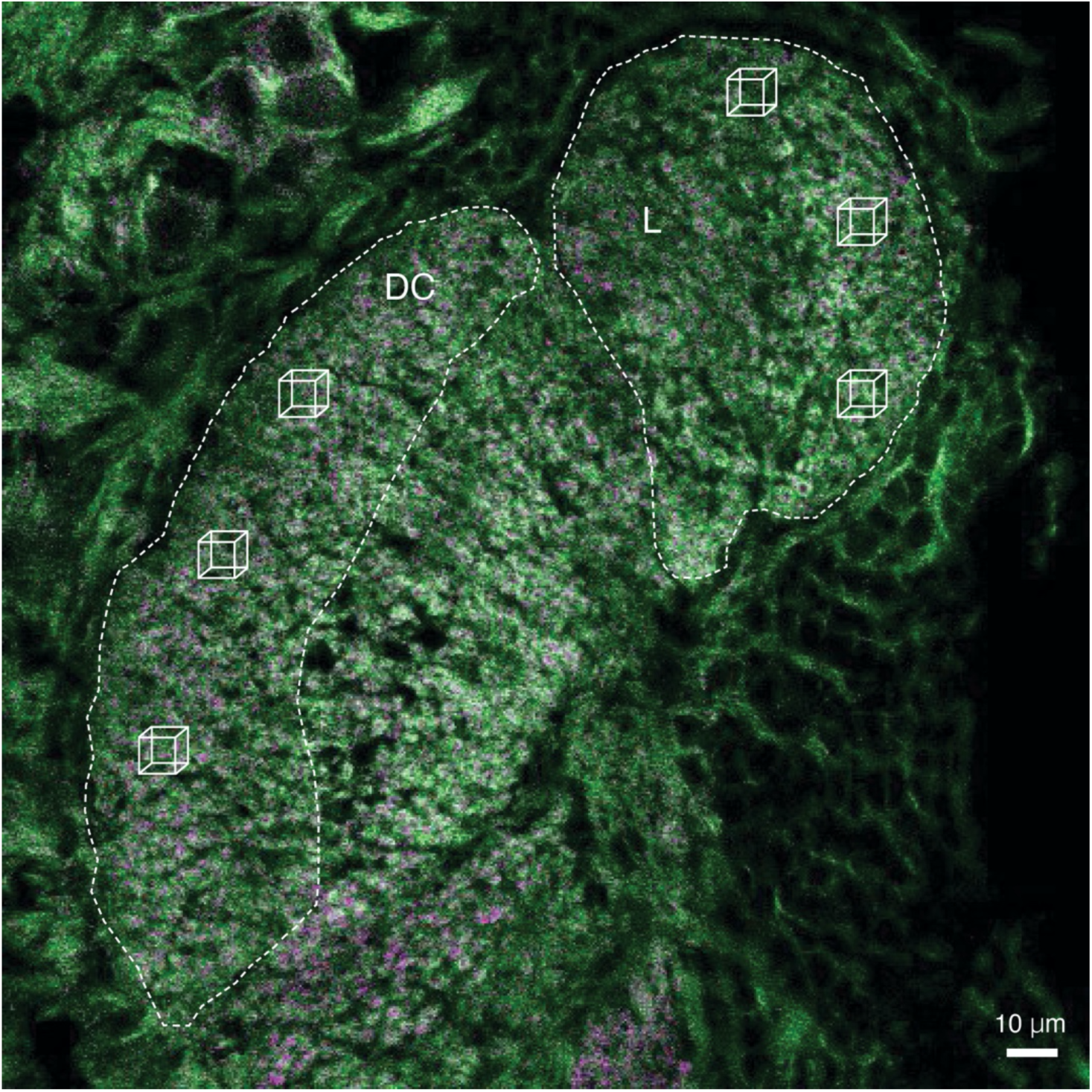
Single optical section of a cryosection of the mushroom body calyx of *Bombus terrestris* stained with ant-synapsin antibody (magenta) and f-actin (green). Dashed lines delineate the two subregions of the calyx analysed (DC, dense collar; L, lip). Three cubes each (8.2 `×` 8.2 `×` 8.2 μm^3^) were randomly placed in the DC and L region to quantify microglomerular density.

### Statistical analysis

Data were analysed using the lme4 and mgcv R-packages (Bates et al. 2014; Wood 2017) in R version 4.1.2 (R Core Team 2021). We performed two main analyses (see below), and within each we analysed MG density in the collar and the lip region separately.

Over the course of the experiment, we recorded foraging efficiency for 2396 foraging trips made by 169 workers. Since not all bees survived until the end of the experiment, we obtained MG density estimates for 65 of these bees, who contributed 1339 foraging trips. During data exploration, we removed one very young bee that was detected as an outlier because it had emerged during (rather than prior to) the foraging period. Consequently, 64 bees contributed to the analysis.

We first explored whether MG density in our sampled bees was predicted by age, size or foraging experience (total time spent outside of the nest during the foraging phase, based on RFID readings). Data exploration revealed a potential non-linear effect of age on MG density for measurements from the lip region but not from the collar region. We thus used a Generalised Additive Mixed Model (GAMM) to analyse the data for the lip region, and a Linear Mixed Model (LMM) for the collar region. In both cases, the response variable was log transformed to reduce the skew of the data, continuous predictors were scaled, and “Colony” was modelled as a random effect. We established the importance of each predictor (fixed factor) by removing it from the full model and evaluating the change in AIC relative to the full model. A predictor was retained if the full model showed improved fit compared to the simpler model (ΔAIC > 2). Interactions between predictors were not included in the model to avoid over-parametrisation considering the sample size.

We then tested the *a priori* hypothesis that MG density predicts foraging efficiency. Because we expected foraging efficiency to increase with foraging experience (Pull et al. 2022), and we had repeated measures of foraging efficiency for each bee, we created an initial LMM with foraging efficiency (mg/min) as the response variable. The fixed effects were: (1) age on sampling (2) size (3) Julian date of initial release and (4) foraging experience (number of days since first foraging bout). Because each bee performed multiple foraging bouts, bee identity was included as a random intercept, nested within colony. Foraging efficiency (n = 64) was transformed to improve model fit using ordered quantile normalisation with the BestNormalize function (Peterson and Peterson 2020). We then tested whether adding MG density (collar or lip) improved the model, based on the change in AIC value. As above, we assessed the importance of a predictor based on the change in AIC value achieved by adding (MG density) or removing (all other predictors) fixed factors from the model.

## Results

### Predicting MG density

For the collar region of the mushroom bodies, we found that neither age, size, total foraging experience nor Julian date of release predicted MG density. In each case, the full model (which contained all predictors) did not perform significantly better than simpler models that excluded the single predictors (ΔAIC compared to full model < 2 in all cases, Table 1a). For the lip region, we identified a non-linear relationship between age at sampling and MG density (Fig 3). A non-linear GAMM thus provided a better fit to the data than a linear model (ΔAIC = 39.11), and the model containing all predictors performed better than a simpler but otherwise identical model that did not contain age (ΔAIC = 5.39). For all the other predictors, the full model showed no improvement over simpler alternatives that did not contain the predictor of interest (ΔAIC < 2 in all cases; Table 1b)

**Figure 3:**
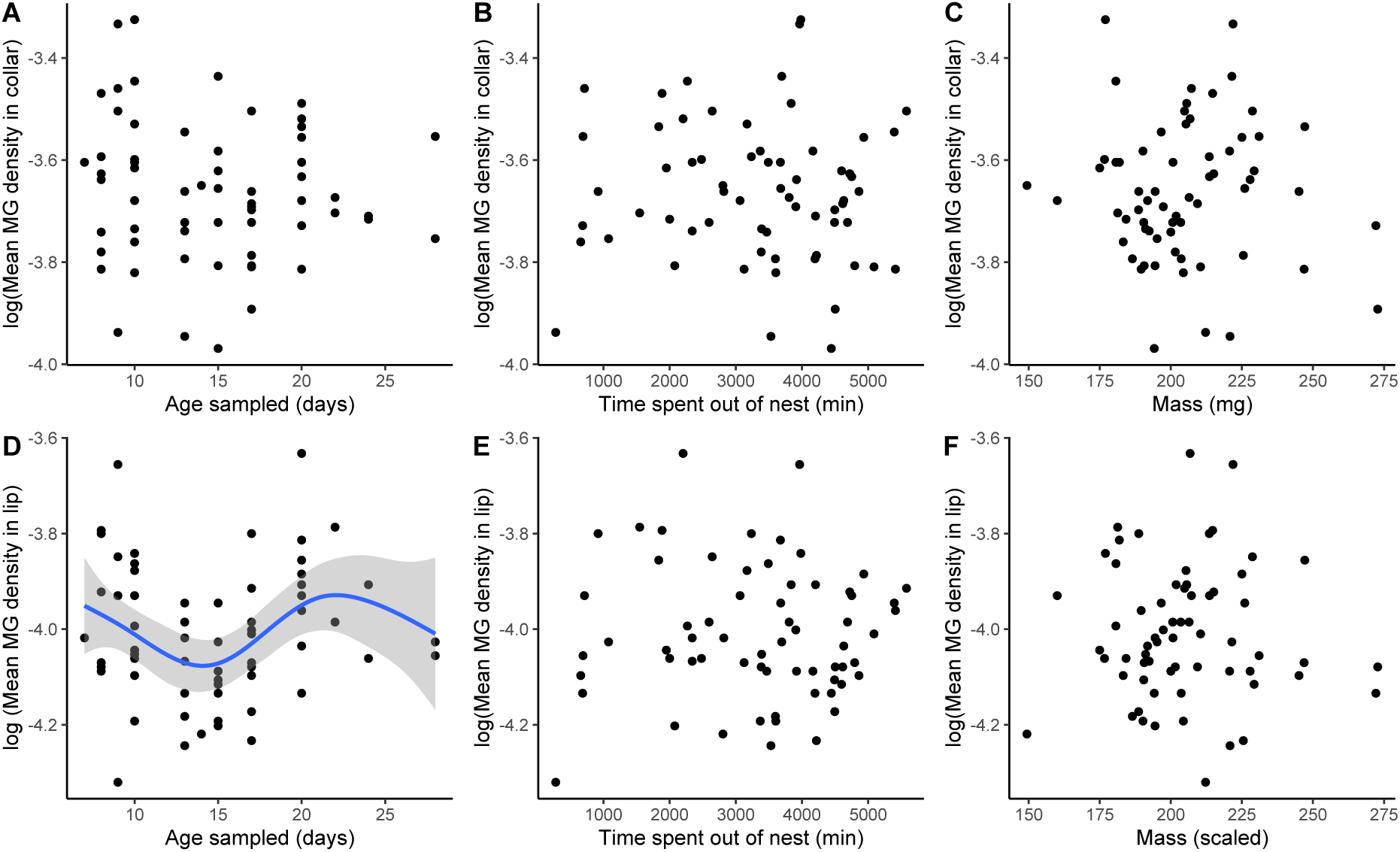
Exploratory analysis of MG density (MG/um^3^) in the (A-C) dense collar and (D-E) lip region of the mushroom body calyx of *Bombus terrestris* workers that had experienced one week of foraging in the wild, immediately before sampling, as a function of age at sampling (A,D), the amount of time that a forager had spent outside of the nest (as measured by RFID readings; B,E) and body mass (C,F). Blue line with 95% CI (grey band) in D indicates non-linear relationship between age at sampling and MG density in the lip region.

**Table 1a:**
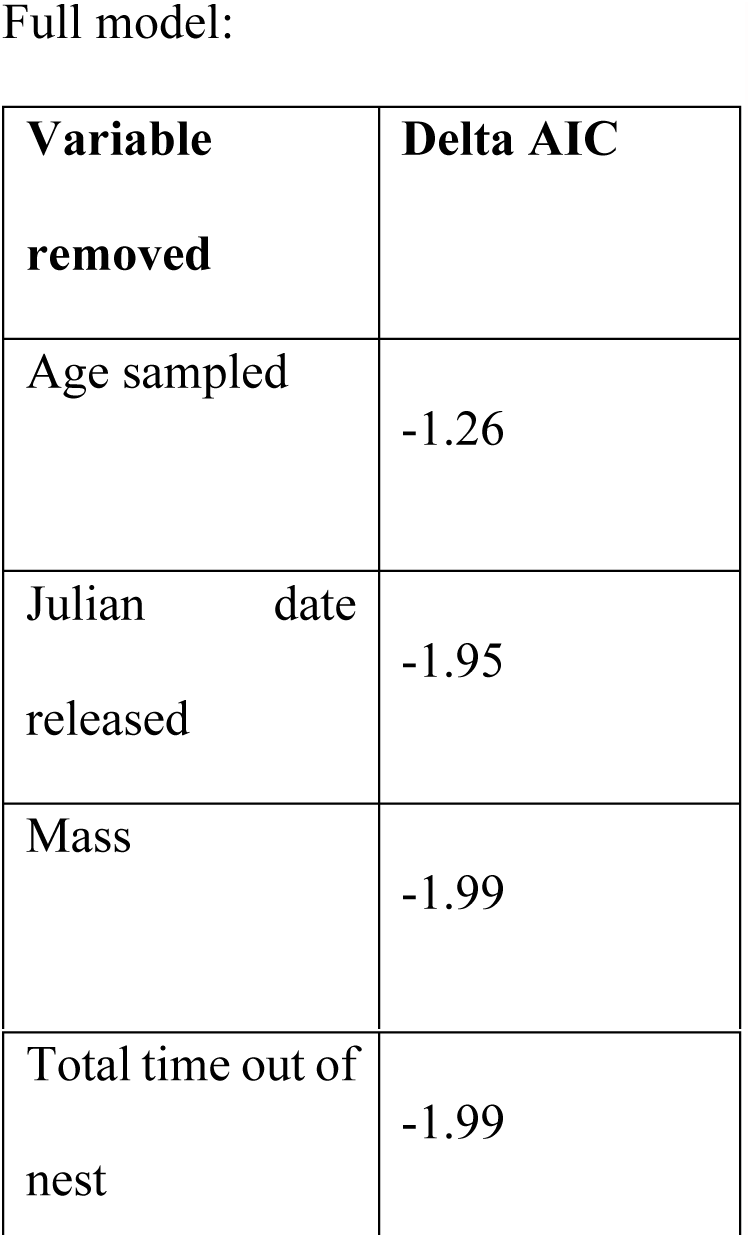
Response variable: MG density in the collar region. Model type: LMER Full model: log(Density in collar) ∼ Age sampled + Julian date released + Mass + Total time out of nest + (1|Colony) All continuous variables are scaled

**Table 1b:**
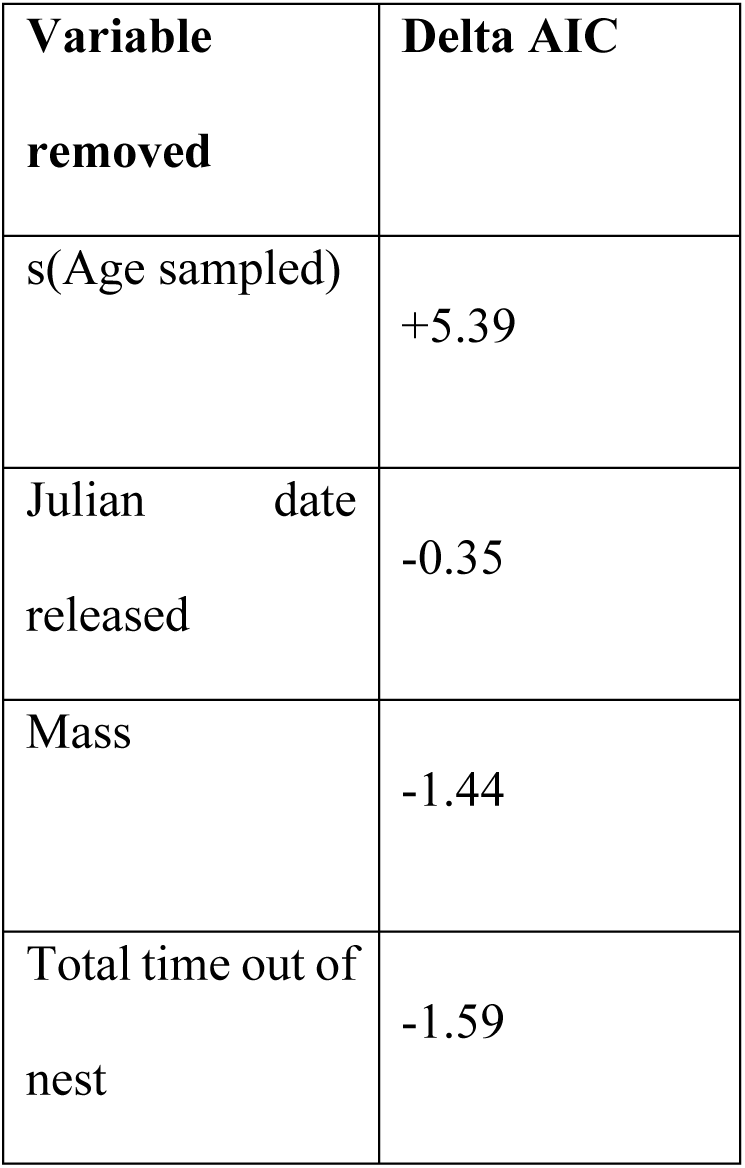
Response variable: MG density in the lip region. Model type: GAMM Full model: log(Density in collar) ∼ s(Age sampled)* + Julian date released+Mass + Total time out of nest + s(Colony, type = “re”)** All continuous variables are scaled *s() = modelled as a non-linear smoother ** s(type = “re”) indicates modelled as a random effect within the GAMM framework

### Predicting foraging efficiency

Individual foraging efficiency increased as bees became more experienced foragers: the full model performed better than a simpler alternative where “days since first foraging trip” was removed (ΔAIC = 76.16, Figure 4). Neither age at release, body size, nor date of release had an effect on foraging efficiency (delta AIC < 2 in all cases, Table 2). We found no evidence to support our *a priori* hypothesis that bees with higher sampled MG density may forage more efficiently, since adding MG density to a model containing all other predictors did not improve fit (ΔAIC < 2 for both lip and collar, Figure 4).

**Figure 4:**
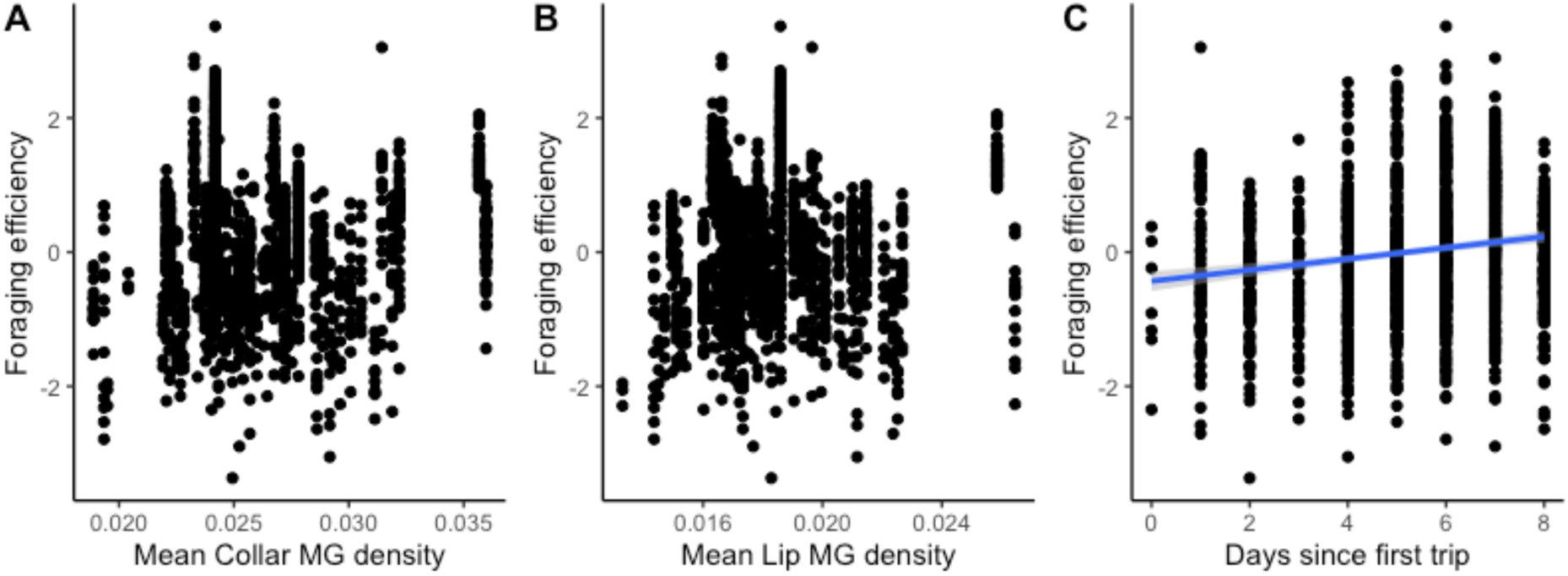
Foraging efficiency (transformed using the function BestNormalize) of 64 workers, over multiple trips per bee, in relation to (A) mean MG density (MG/um^3^) in the dense collar region; (B) mean MG density (MG/um^3^) in the lip region; (C) foraging experience (note that including a quadratic term did not improve the model fit).

**Table 2:** Response variable: foraging efficiency. Model: Foraging efficiency* ∼ Age + Julian date released + Mass + Days since first trip + (1|Colony/ID) * transformed using ordered quantile normalisation with the BestNormalize function (Peterson and Peterson 2020)

| Variable | Removed or added? | Delta AIC |
| --- | --- | --- |
| Age | Removed | -0.82 |
| Julian Date Released | Removed | -1.89 |
| Mass | Removed | -1.98 |
| Days since first trip | Removed | +76.16 |
| Microglomerulus Density (Collar) | Added | +1.96 |
| Microglomerulus Density (Lip) | Added | +1.117 |

## Discussion

Our results did not reveal any association between MG density and foraging efficiency in bumblebees. Previous work has shown that MG density in the MBs changes in response to learning events (Hourcade et al. 2010; Li et al. 2017), predicts learning ability (Li et al. 2017; Cabirol et al. 2018), and coincides with the onset of foraging (Muenz et al. 2015; Kraft et al. 2019). However, our results suggest that these changes, if they are related to learning ability or the experience of learning, are not reflected in improved foraging efficiency in bumblebees, at least within the confines of our protocol. Instead, we found that the number of days between a focal foraging trip and the first outing of a bee was the only predictor of foraging efficiency within our test cohort. In other words, in line with previous findings (Pull et al. 2022) foraging efficiency improved with experience, as bees learnt about their environment.

Our analysis identified a potential non-linear effect of age on MG density in the lip region of the mushroom bodies, whereby MG density decreased until ∼15 days of age, but returned to similar levels in older bees. Previous work also suggests that MG density is high on emergence and decreases quickly (Kraft et al. 2019), although note that our study did not include very young bees at an equivalent stage, because all sampled individuals had foraged. We are cautious in any interpretation because this non-linear relationship was identified *post hoc,* and the pattern should be further explored through an inference-based approach (Tredennick et al. 2021). Although MG density in honeybees follows a similar pattern with age, initially decreasing at the start of foraging followed by a subsequent increase (Cabirol et al. 2018), MB plasticity in honeybees is to some extent driven by the onset of foraging rather than aging itself (Ismail et al. 2006); age-based polyethism means that the two variables are correlated in this species. In bumblebees, we found no evidence that foraging experience – which was independent of age in our study – predicted MG density. The findings from our study and others (Pull et al. 2022), suggest that MG density may be fairly robust to the influence of exposure to the visual and olfactory cues of the real-world foraging environment in bumblebees.

Our experimental setup allowed us to directly test the relationship between foraging efficiency, as the fitness-relevant potential product of learning, and MG density. We hypothesized that if MG density either responds to (Hourcade et al. 2010; Li et al. 2017) or promotes (Fahrbach and Van Nest 2016; Li et al. 2017; Cabirol et al. 2018) learning about the environment, variation in MG density might predict variation in foraging efficiency, but this was not the case. However, a recent study by Pull et al. (2022) that used an experimental setup and location almost identical to ours found that the relationship between a cognitive ability (specifically a short-term form of memory, as assayed through a radial arm maze) and foraging efficiency is not straightforward, and can vary across the year. We tested our bees in the height of the UK summer, when food availability is low (Pull et al. 2022) and performance in an associative learning task has been previously shown to predict foraging efficiency, at least at the colony level (Raine and Chittka 2008). However, future studies could fruitfully explore how the relationship between learning ability and foraging efficiency changes depending on ecological conditions, and particularly whether MG density varies between bees exposed to complex, rich spring environments *vs* those foraging in the summer dearth.

Another explanation for our results could be that cognitive abilities play little role in foraging under harsher ecological conditions where food is scarce and exploratory activity would be of greater importance (Pasquier and Grüter 2016). Fidelity to a route, which requires learning, decreases the likelihood of discovering a new, more profitable food source by chance, and might conceivably be detrimental in a poor and changing environment, although this has not been tested. Furthermore, a recent study failed to find neural correlates between MB extrinsic neurons’ activity and exploratory behaviour in bumblebees (Jin et al. 2020) which suggest exploratory behaviour is unlikely to produce the repetitive stimulation necessary to cause an increase in synaptic density. Nevertheless, exploration is expected to lead to substantial energetic costs that colonies might find harder to balance in a poorer environment. Similarly, increased cognitive abilities also come at a cost and have been shown to trade-off with survival (Mery and Kawecki 2005). More research is therefore needed to determine the trade-offs involved with exploratory activity alone and comparison with cognitive abilities. Overall, our experiment highlights the gaps in knowledge on foraging and its relationship with cognition and neuroanatomy.

